# Effective spatio-temporal regimes for wound treatment by way of macrophage polarization: A mathematical model

**DOI:** 10.1101/2021.03.25.436953

**Authors:** Jiahao Xue, Ksenia Zlobina, Marcella Gomez

## Abstract

Wound healing consists of a sequence of biological processes often grouped into different stages. Interventions applied to accelerate normal wound healing must take into consideration timing with respect to wound healing stages in order to maximize treatment effectiveness. Macrophage polarization from M1 to M2 represents a transition from the inflammatory to proliferation stage of wound healing. Accelerating this transition may be an effective way to accelerate wound healing; however, it must be induced at the appropriate time. We search for an optimal spatio-temporal regime to apply wound healing treatment in a mathematical model of wound healing. In this work we show that to maximize effectiveness, treatment must not be applied too early or too late. We also show that effective spatial distribution of treatment depends on the heterogenity of the wound surface. In conclusion, this research provides a possible optimal regime of therapy that focuses on macrophage activity and a hypothesis of treatment outcome to be tested experimentally in future. Finding best regimes for treatment application is a first step towards development of intelligent algorithms of wound treatment that minimize healing time.

## Introduction

Wound healing is an important healthcare problem, and by far there is no decisive finding for the best therapy [1]. The best treatment is one that can heal patients as fast as possible. However, the treatment may also exert no effect or even a negative effect on healing tissues if not applied appropriately [2]. For example, many treatments are suggested to accelerate certain phases of inflammation but have little effect on other phases. In some cases, treatment induces toxic side-effects [3]. For this reason each treatment should be applied only at appropriate stages of wound healing. Finding the best regimes for treatment application is a first step towards development of intelligent algorithms for wound treatment that minimize healing time.

Mathematical modeling has yielded considerable understanding of wound healing, particularly the aspects that are difficult to investigate empirically [4] [5]. Macrophage polarization modeling, to our knowledge, has been applied to several biological situations but not to wound healing [6] [7]. In this study, we investigate mathematically the regimes of wound treatment by an abstract actuator that accelerates macrophage polarization. This type of treatment can have adverse effects if applied improperly [8] [9]. For example, a fast transition from the inflammatory to reparative stage may slow down cleaning of the wound of debris (damaged cells and infection). We focus our attention on healthy wound healing, with the objective of shortening the time to wound closure. To this end, we create a simple model with only one stable state – the healthy one. We note that our model does not capture switching between chronic and normal wound healing regimes. However, in order to capture trade-offs of early treatment, we examine wound cleaning time as well, with the understanding that any remaining wound debris can be an indicator of prolonged inflammation and potential infection preventing wound closure. In summary, we examine wound healing time and wound debris cleaning time in response to different spatio-temporal signals inducing macrophage polarization. Overall, decoupling the different modalities of wound healing trajectories reduces complexity of the model and allows us to gain intuition for optimal treatment strategies. We find that actuation of M1 to M2 polarization must be applied with care and optimal timing can depend on the duration of the treatment, time of initiation, placement of the actuator and initial distribution of wound debris.

### Macrophages in wound healing

Wound healing is usually divided into four stages: hemostasis, inflammation, proliferation and reparation [10]. During hemostasis, a blood clot is formed to stop bleeding [11]. At the inflammation stage leukocytes arrive and clean the wound of debris and infection. During proliferation, a temporary extracellular matrix is produced and the microenvironment serves to produce sufficient amount of healthy tissue cells [12]. Finally, in the last stage known as reparation, new tissue is restored.

Macrophages are important players of inflammation and proliferation stages in wound healing. Initially macrophages appear in the wound as pro-inflammatory or classically activated macrophages M1. They fight against infection and debris mainly by phagocytosis [13]. By that time, neutrophils have completed their function and begin to die by apoptosis. M1 macrophages can phagocyte apoptotic neutrophils in a process called efferocytosis and become anti-inflammatory or alternatively activated macrophages M2 [14] [15].

M1 macrophages are important to eliminate pathogens and other small particles, such as engulfing and absorbing pathogens and secreting pro-inflammatory cytokines. However, too much M1 can also lead to tissue harm and chronic inflammation [16]. On the contrary, M2 macrophages are associated with the subsiding of inflammation and transition to the proliferation stage of wound healing [17] [18]. Although M2 is able to diminish inflammation, too much M2 can cause some negative effects such as allergies for example [19].

The terms M1 and M2 were coined following an in vitro culture, in which particular molecules were found to stimulate macrophages to transform to M1 and M2 [20]. Several surface molecules are assumed to be markers of M1 and M2 subtypes. Later studies of macrophages revealed more subtypes like M2a,b,c [21]. In practice, M1 and M2 have overlapping functions, i.e., many functions are shared by macrophages of different subtypes, thus making their research more difficult.

In our approach we assume that M1 macrophages are responsible for cleaning the wound of debris and M2 macrophages for inducing proliferation processes. In this model they don’t share functions. We divide macrophages by function rather than by surface markers.

## Mathematical model

### Mathematical model of wound healing

Consider a wound of radius R. Let the axis x to be directed from the wound center to the edge (Fig. 1a). Concentrations of substances and populations of cells are functions of x. A schematic of modeled biological processes including macrophages participation in the wound healing is shown in Fig.1b.

**Fig 1.**
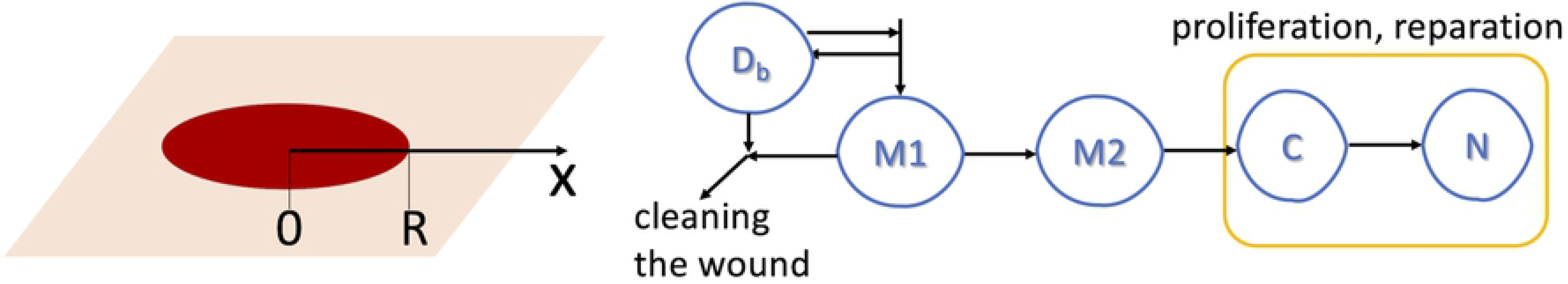
Geometry and scheme of model. (a) Geometry of the model: x-axis is directed from wound center; R is the wound radius. (b) Scheme of the model of wound healing. Wound debris (*D*_*b*_) attracts macrophages M1 that remove debris. Macrophages M1 become M2. Macrophages M2 induce production of temporary tissue (C) that helps new tissue (N) to grow.

*D*_*b*_ is wound debris consisting of damaged cells and bacterial cells promoting infection. We assume the wound debris to be non-active and only eliminated by M1 macrophages:

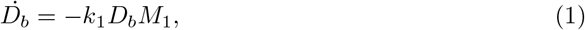

*M*_1_ is the population of M1 macrophages, which are attracted by debris (*k*_2_*D*_*b*_) and removed in the reactions of debris elimination (*k*_1_*D*_*b*_*M*_1_), macrophages polarization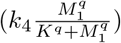 and natural death (*k*_*d*1_*M*_1_). Spatial migration of macrophages is described by the last term in the equation:

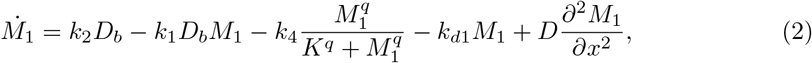

*M*_2_ is the population of M2 macrophages whose dynamics are driven by M1 polarization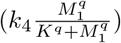, death rate (−*k*_*d*2_*M*_2_) and migration as follows:

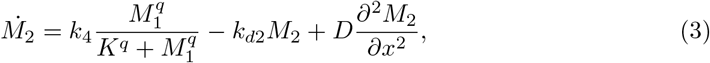

*C* is temporary tissue – sum of macromolecules and enzymes of extracellular matrix that help new tissue to develop. This variable appears as a result of M2 action and disappears after remodeling:

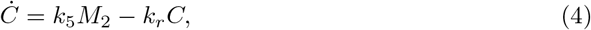

The state *C* is an intermediate state leading to the growth of new healthy tissue, N. To model growth of new healthy tissue, we assume that the top layer of the skin, epithelium, grows as a sheet from the edge of the wound [22]. The behavior of new tissue growth can be described by equations similar to classical models of a running wave [23]. However, this is possible only in the presence of temporary tissue, so the rate of new tissue growth is proportional to *C*:

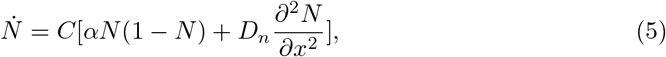

In order to reduce the number of parameters, we rewrite the system in non-dimensional form by introducing new variables:

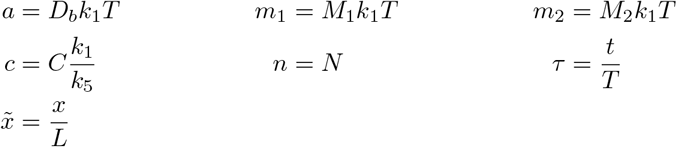

where *T* and *L* are characteristic time and length scales. The system of equations in non-dimensional form is

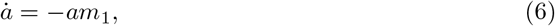

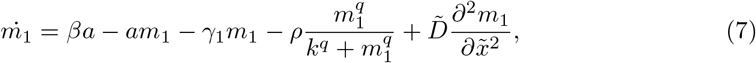

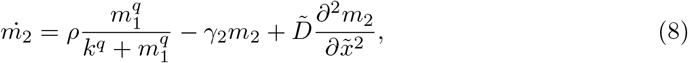

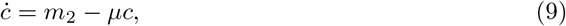

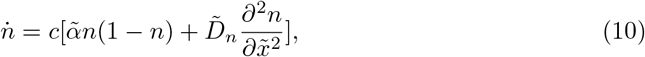

where *β* = *k*_2_*T*, *ρ* = *k*_1_*k*_4_*T*^2^, *γ*_1_ = *k*_*d*1_*T*, *γ*_2_ = *k*_*d*2_*T*, *μ* = *k*_*r*_*T*, 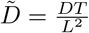, 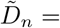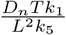, 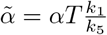, *k* = *Kk*_1_*T*.

The values of parameters used in numerical simulations are listed in Table 1.

**Table 1.** The values of parameters used in numerical simulations.

| Parameter | Value | Reference | Parameter | Value | Reference |
| --- | --- | --- | --- | --- | --- |
| R | 3 mm | | $\delta$ | 8 hours | |
| T | 3 days | | $\gamma_1$ | 0.1 | [26] [27] |
| L | 0.03 mm | | $\gamma_2$ | 0.1 | |
| $\beta$ | 1 | [24] [25] | $\mu$ | 0.2 | C disappears in 3 weeks |
| $\rho$ | 0.1 | [28] | $\tilde{D}$ | 0.32 | |
| k | 0.05 | | $\tilde{D}_n$ | 0.0003 | unpublished data from UCD |
| q | 5 | | $\tilde{\alpha}$ | 1.8 | |

The Initial conditions are:

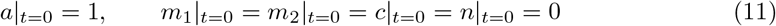

We assume zero-flux boundary conditions for macrophages on the right and left boundaries of the considered region:

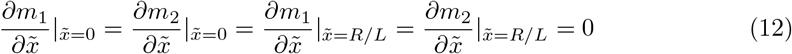

New tissue is assumed to be constant on the edge of the wound and non-moving through the center of the wound:

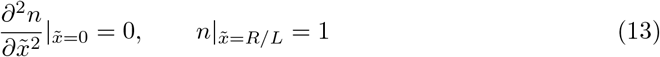

The results of wound healing model simulations are shown in Fig.2. Wound radius is measured as the distance from wound center to the location 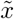 where 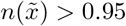

**Fig 2.**
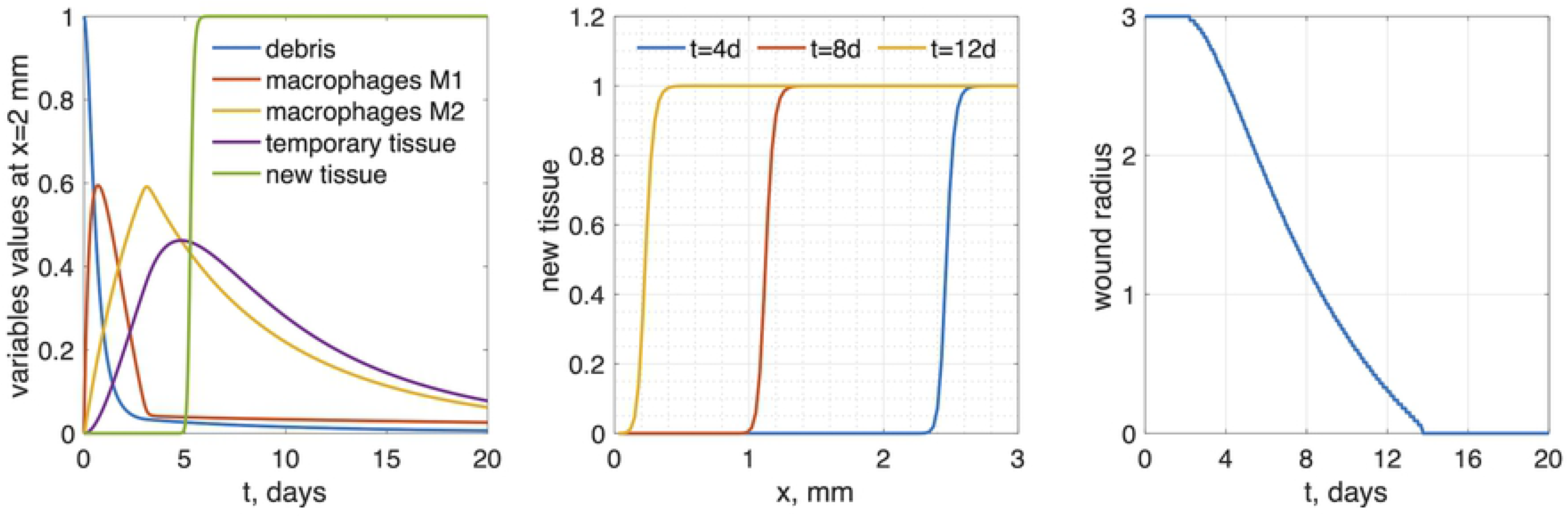
Results of wound healing model simulations, R=3mm. (a) time-dependence of all variables at distance x = 2 mm from wound center (b) new tissue profiles as functions of x for several time points. (c) wound radius vs time: wound healing time is 13.77 days.

### Model of wound with actuator

In order to investigate regimes of wound healing treatment we include actuator enforced macrophages polarization into the model. This actuator is applied at point *x* = *x*_*p*_. We suppose that the actuator delivers some substance at the point of application and its concentration *θ* is distributed by diffusion. The concentration distribution of the injected treatment in space is assumed to be quasi-stationary (see Fig. 3a):

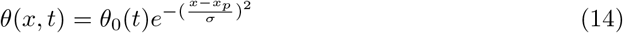

where *θ*_0_ is the amplitude of the treatment controlled by the actuator. One can see that *θ* = *θ*_0_ at the location of actuator, *x* = *x*_*p*_. In this work we assumed *σ* = 0.45 mm. The treatment substance affects macrophage polarization, so equations for *m*_1_ and *m*_2_ with actuators may be rewritten:

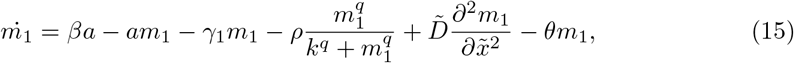

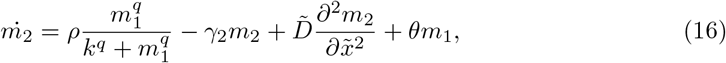

**Fig 3.**
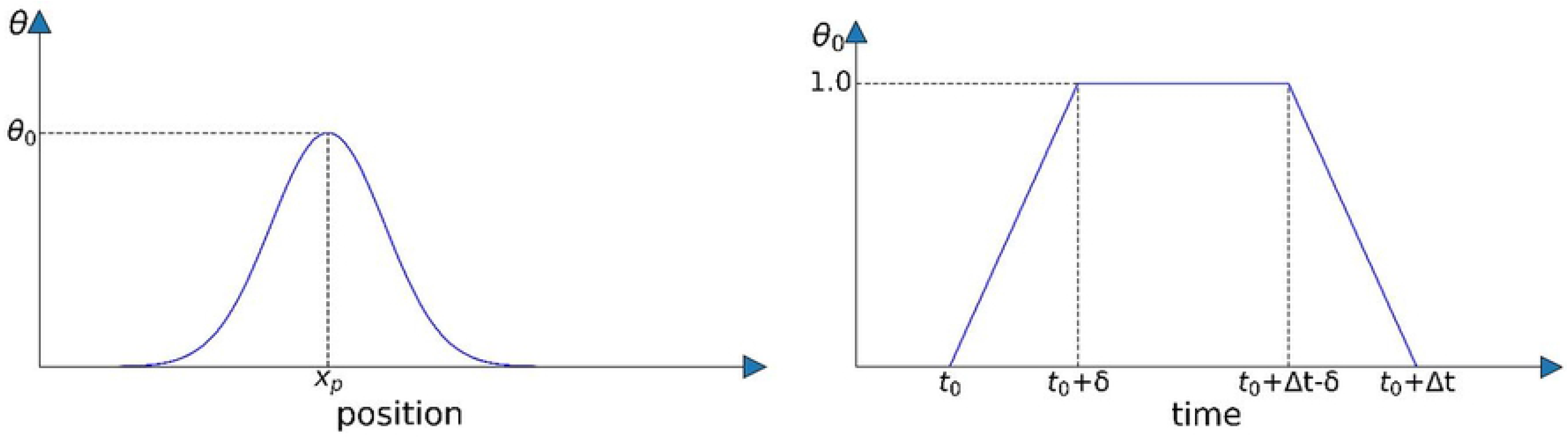
Spatio-temporal characteristics of treatment induced by actuator. (a) Treatment substance distribution in space. The actuator is located at *x* = *x*_*p*_. (b) Time dependence of the actuator amplitude.

*θ*_0_(*t*) is the time-dependent regime of treatment controlled by an actuator. In order to find the best regime, we test the model with impulses of actuator treatment of duration Δt beginning at time *t*_0_ (see Fig. 3b):

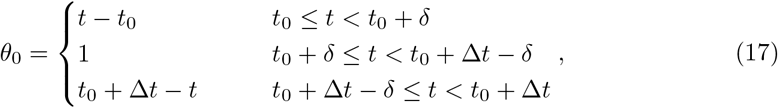

We tested other impulse-like shapes of time-dependence of the actuator and general results are the same. We present the piece-wise linear function for simplicity. In order to make actuator signal switching on and off smoother, we assume linear growth and decrease in the beginning and end of the signal.

## Results

We define wound healing time as time from injury (t=0) to the moment when wound radius reaches zero. Application of the actuator accelerating macrophages polarization in the model decreases the time of wound healing. The results are shown in Fig.4–6.

**Fig 4.**
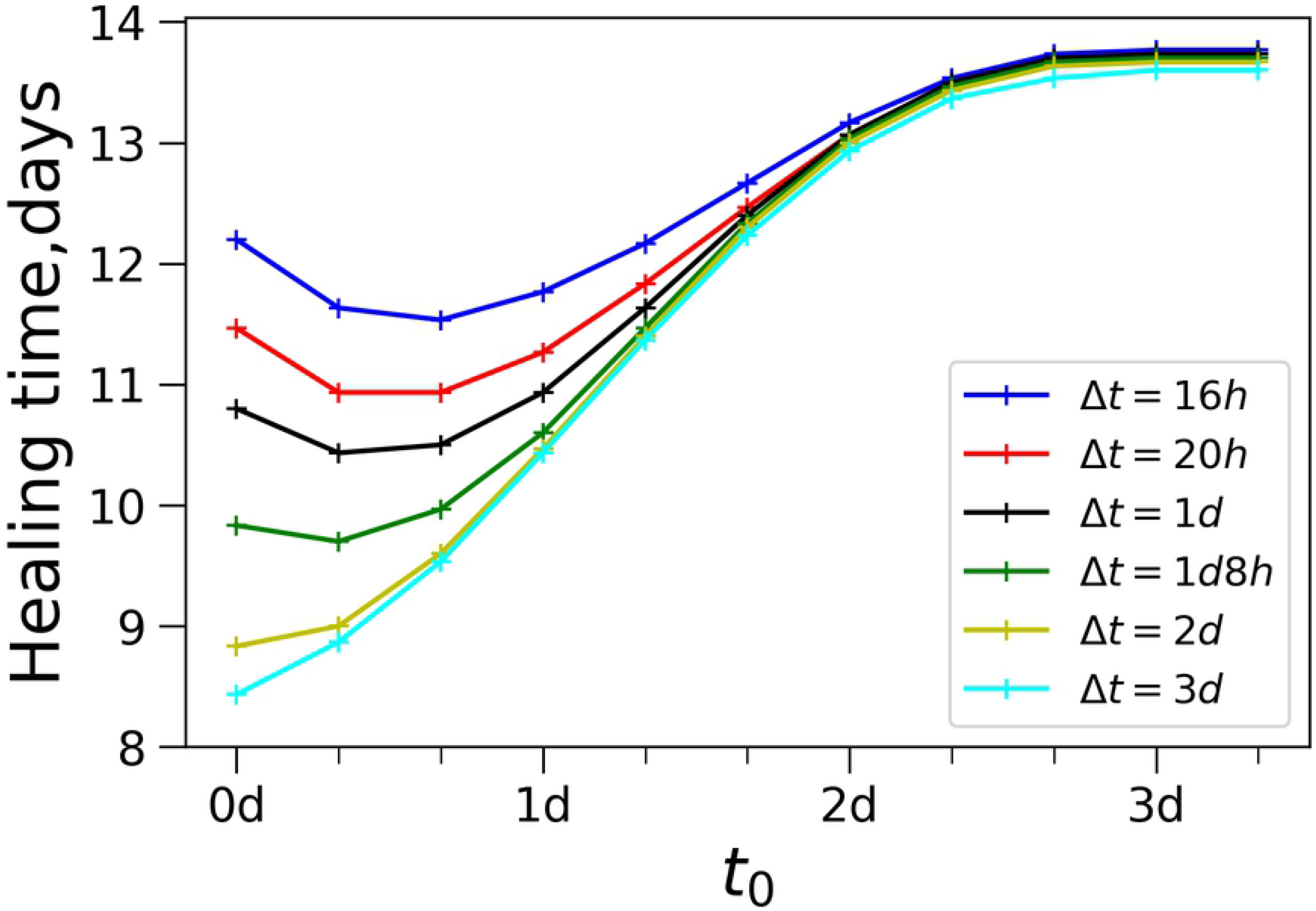
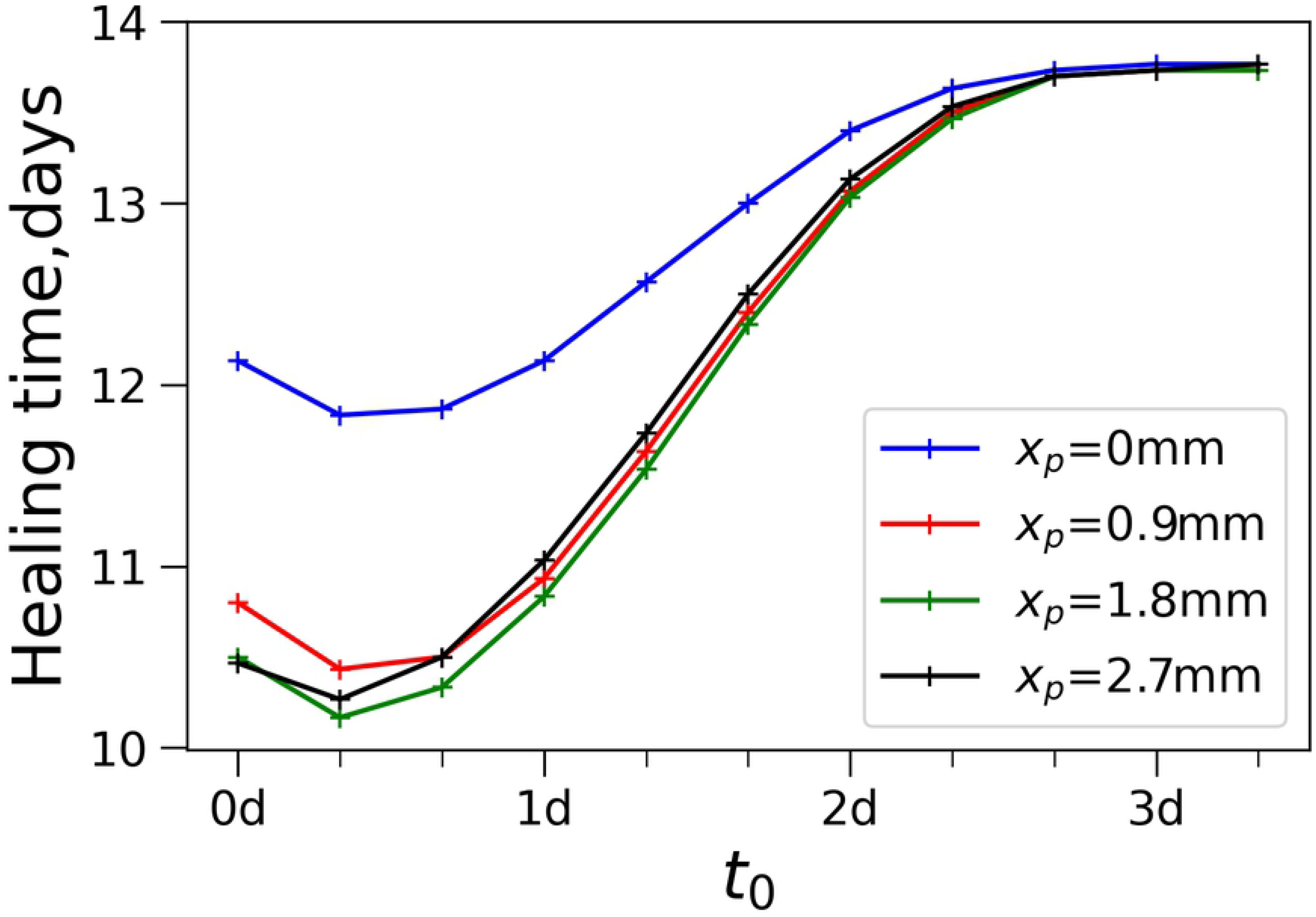
Wound healing time dependence on the treatment beginning time *t*_0_. (a) plots for different treatment durations Δ*t*, *x*_*p*_ = 0.9 mm. (b) plots for different actuator position *x*_*p*_, Δ*t* = 1 day

Beginning time of the actuator plays an important role in wound healing (see Fig.4). Large values of *t*_0_ make treatment less effective: wound healing time increases as *t*_0_ increases. In our simulations for *t*_0_ > 3d the treatment doesn’t have any effect: the value of healing time tends to the value of healing without treatment (13.77 days for given set of parameters).

Interestingly, for shorter treatment durations Δt, healing time plots have a minimum away from extremes(see plots for Δ*t* < 2d in Fig. 4a). This means that there is a non-trivial optimal treatment beginning time *t*_0_. In other words, there is a short window of time during wound healing when artificial acceleration of macrophages polarization is most effective. The plots for other actuator positions *x*_*p*_ are shown in the S1 Fig. Similar trends are observed for different placement of the actuators.

Fig.5 shows how wound healing time depends on the duration of treatment. The longer the treatment time Δt is, the better its effectiveness at accelerating wound healing is. However, for Δt larger than 2-3 days wound healing time approaches a lower bound. This means that further prolongation of the treatment has minimal effect. The plots for other values of *x*_*p*_ are shown in S2 Fig. Again, we find a similar trend regardless of actuator placement.

**Fig 5.**
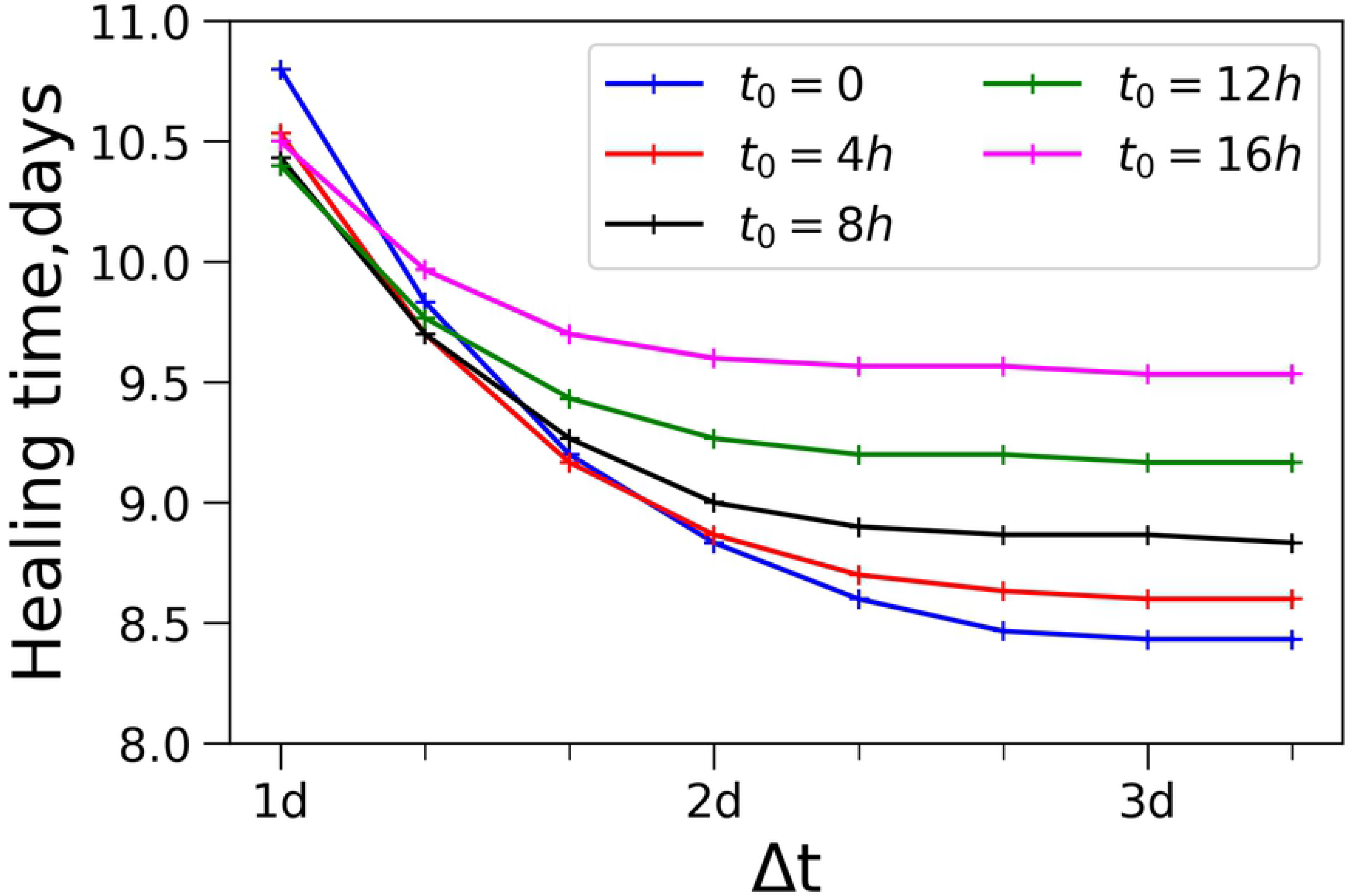
Wound healing time dependence of treatment duration Δt for different beginning time *t*_0_. *x*_*p*_ = 0.9 mm.

Fig.6a shows the dependence of the wound healing time on the actuator position *x*_*p*_. The treatment substance in this model is approximated by a Gaussian function on a bounded domain, so the integral of this function depends on the position of maximum, *x*_*p*_. When the actuator is close to the edge of the considered region the integral amount of treatment becomes smaller, as if the treatment substance would go outside the considered region. This explains why treatment effect is partially lost for *x*_*p*_ close to 0 and 3 mm. Besides this boundary effect, the plots are slightly decreasing from wound center to the edge. This implies that an actuator located close to the wound edge is beneficial. The plots for other Δ*t* and *t*_0_ are shown in S3 and S4 Figs.

**Fig 6.**
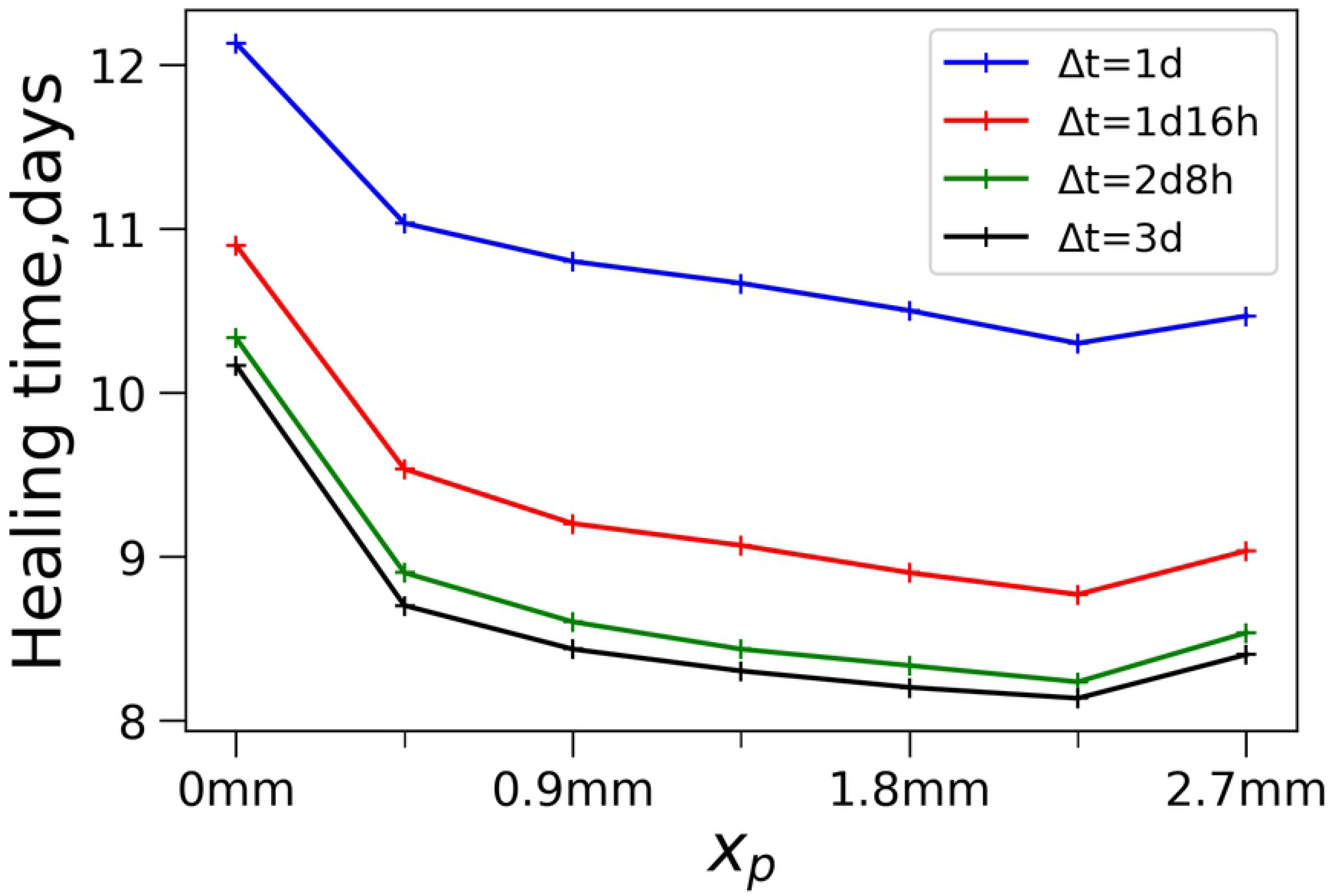
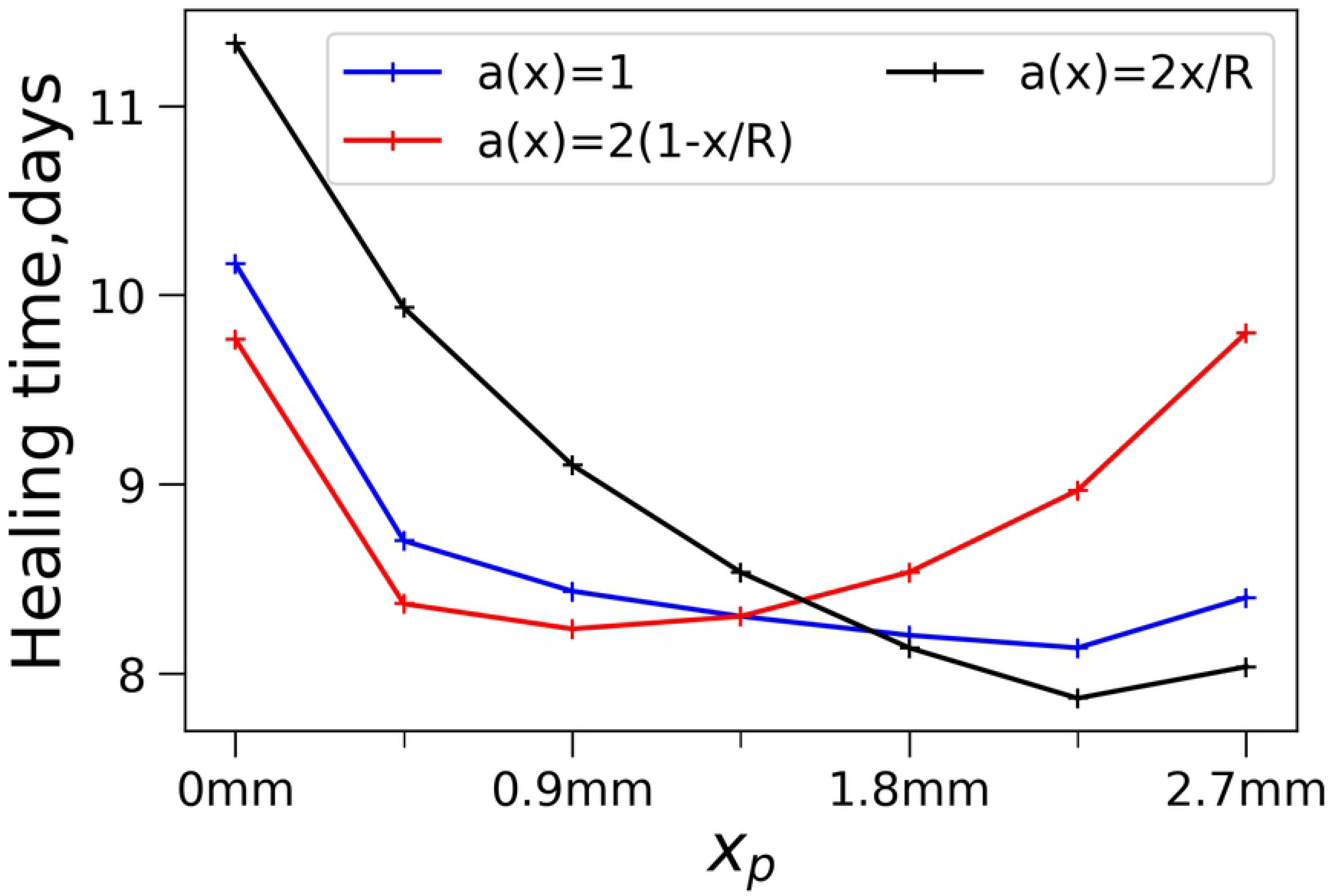
Wound healing time dependence on the actuator position *x*_*p*_. (a) plots for different treatment duration Δ*t*, *t*_0_ = 0*h* (b) plots for different initial distribution of debris in the wound, *t*_0_ = 0*h* Δ*t* = 3days

However, this might be the consequence of the initial uniform distribution of the debris in the wound (see initial conditions). We made additional simulations with debris accumulation in the center and at the edge of the wound. Alternative initial conditions for the debris variable take the following form:

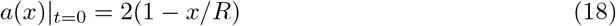

and

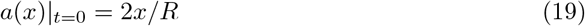

Wound healing time dependence on *x*_*p*_ for the three different types of initial conditions on debris distribution is shown in Fig.6b. One can see that both debris amount and its distribution in the wound bed affect the dependence of healing time on actuator placement.

For the case *2* (1-x/R) (more debris in the wound center), the actuator should be placed near the center. For the case *a*(x)=2x/R, there is more debris on the edge. Correspondingly, if the actuator is placed near the edge, we can get much shorter healing times. One can see that the best choice of the best position of the actuator is sensitive to the distribution of the debris in the wound. The best position cannot be estimated in the framework of this rough modelling and must be investigated experimentally.

Fig.7 demonstrates the limitations of treatment regimes. In addition to wound healing time, we define the time of wound cleaning of the debris. Because debris is diminishing variable in our model, it tends to zero as time goes to infinity. We define wound cleaning time as the time when the maximal value of debris across the wound bed falls below a small threshold 0.04:

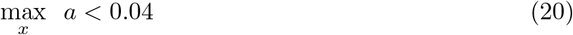

**Fig 7.**
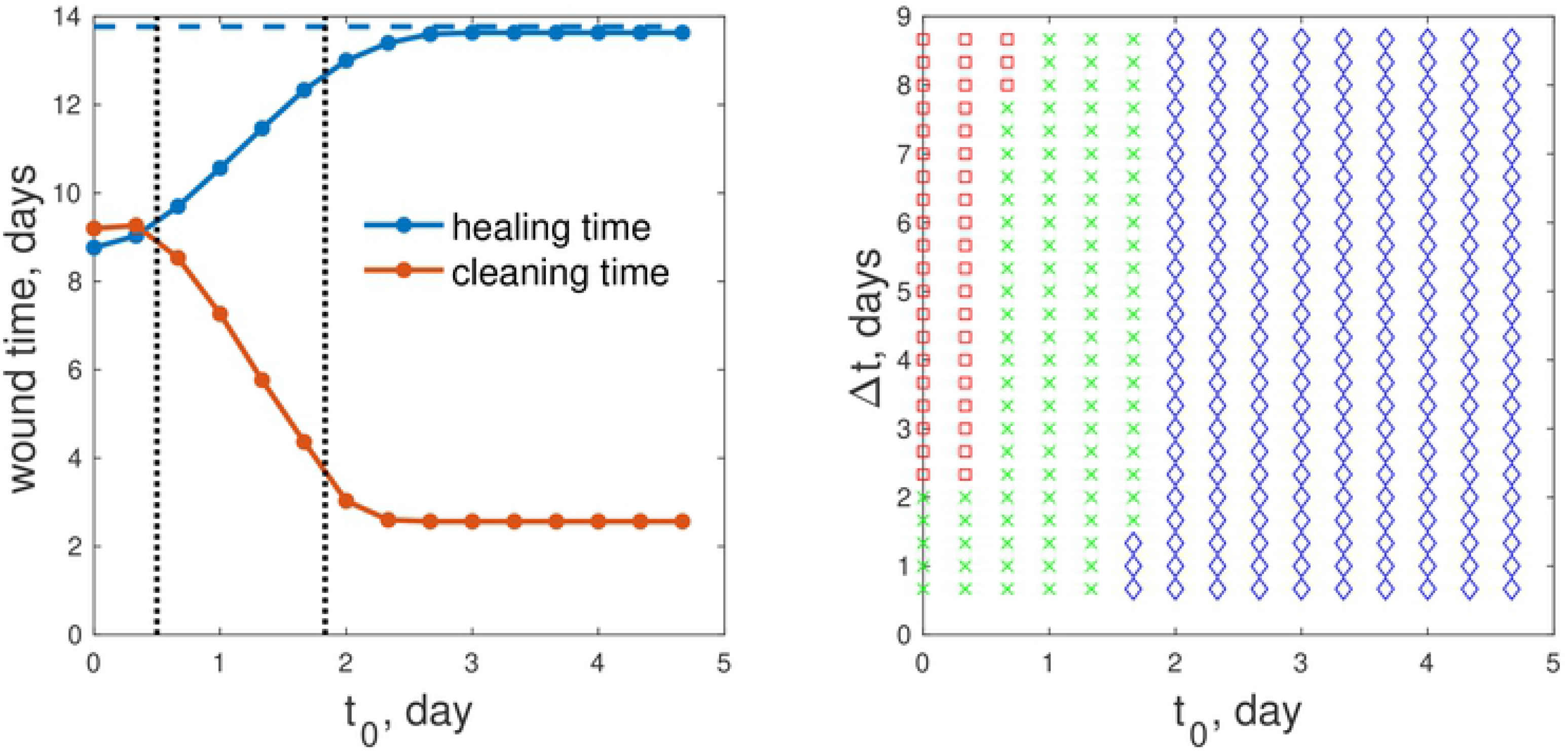
Optimal regimes of wound treatment. (a) wound healing time and wound cleaning time as functions of treatment beginning time, *t*_0_ (Δ*t*=2d8h). Blue dashed line: healing time without treatment. (b) Parametric plane for the choice of the best treatment regime: green crosses – the best regimes of treatment; red squares – wound cleaning takes longer time than wound healing; blue diamonds – wound healing time diminishes less than 10% in comparison with non-treated wound (*x*_*p*_=0.6 mm).

Because the treatment applied in this model accelerates M1 to M2 transition, it not only accelerates the reparation stage but removes M1 cells performing debris removal. Too large application of this treatment can make debris removal too slow —this is the cost of accelerating the reparation stage. Fig.7a shows that the smaller the wound healing time is, the larger the time of wound cleaning of debris is. For some scenarios, wound cleaning may take even longer than wound closure, which is physically unrealizable and indicative of complications in wound healing.

We can see two main limitations of the treatment regimes, divided by vertical dotted lines in Fig 7a. The regimes with small *t*_0_ (left of the first dotted line) lead to too slow wound cleaning. Regimes with very large *t*_0_ (right of second dotted line) provide almost no benefit in wound healing time. That is, the difference between wound healing time with treatment and without treatment is less than 10%. In the case the potential side effects of applying the treatment may overweight any benefit.

Only the regimes with middle values of *t*_0_, between vertical dotted lines in Fig7a, make wound healing time smaller and wound cleaning time not too large.

Fig.7b shows these 3 types of regimes in the (*t*_0_,Δ*t*) parametric plane. The regimes marked as red squares correspond to regimes when wound cleaning takes longer time than wound healing, whereas blue diamonds correspond to regimes when wound healing time diminishes less than 10% in comparison with non-treated wound. Green crosses represent the effective regimes of wound treatment. Of course, one can choose more stringent conditions for optimal regimes (maybe 10% healing time improvement is not enough).

Among all the regimes presented in Fig 7b the best healing time (9 days) is observed for the treatment regime *t*_0_=0 and Δ*t*=2 days. In practical situation there might be additional constrains for the duration of treatment, or for maximum concentration that is not considered here. However, our model demonstrates the principles of wound treatment regime optimization.

## Discussion

Wound healing is a sequence of stages, with different cells performing different functions. Therefore, it is reasonable to assume that at each stage, different medications should be applied to improve healing. However, to our knowledge, there are not many studies of wound treatment regimen. In this work, we attempted to find the best wound treatment regimen by affecting the polarization of macrophages. Other type of medicaments may be investigated in the same manner.

The presented mathematical model takes into account the presence of two types of macrophages in the wound—M1 and M2. It is believed now that in addition to the two main types, there are several subtypes of macrophages. All types of macrophages were found in vitro as a result of a specific activating stimuli. The exact types of macrophages in wounds have not been established and are considered to be roughly similar to those found in vitro. It is most likely that macrophages of different types can be present in the wound simultaneously. However, the functions that these macrophages perform are more important for the healing process than the markers found in vitro. Therefore, in this model, we clearly divide the functions of the macrophages into inflammatory and reparative, keeping in mind that such pure cell lines may not exist in reality. This separation of functions of macrophages gave us the opportunity to draw up a rather simple naive model and identify general patterns of the effect of the treatment regimen on the wound healing time.

Our results imply that the treatment with activation of macrophage polarization should not take place too early. Otherwise, macrophages M1 have no enough time to eliminate wound debris. This is especially important for the infected wounds when the debris consists not only of damaged cells but of bacterial pathogen [31]. In the case of bacterial pathogen, the first equation in the model should include pathogen reproduction, the regimes with pathogen persistence may occur. This may lead to continuation of M1 recruitment, inflammation persistence and prevent full wound closure. On the other hand, treatment with activation of macrophage polarization should not begin too late, because it becomes not effective and toxic side effects are unknown.

The model demonstrates mathematically why actuating M1-M2 is good only at the appropriate stage. For the particular values of parameters the best actuating stage is between 0.7 and 3.1 days.

This model has its limitations. The model constructed in this work does not take into account the details of the polarization of macrophages due to the poor knowledge of this issue in vivo [26]. It is known that M2 macrophages can appear in response to stimulation by certain cytokines, for example, IL4, produced by basophils and mast cells [18]. There are indications that macrophages M1 are converted to M2 after they phagocyte apoptotic neutrophils [29]. We made a model in which M1 population is replaced by M2 although the mechanisms of this transition in vivo are not well understood.

With modern development of electronics, it is possible to reach not only high temporal but high spatial distribution of the medication application [30]. That’s why we made an attempt to find the best position of the wound treatment. The lack of knowledge about debris distribution in the wound makes it difficult to predict the best actuator position. However, we demonstrate several possible solutions for different debris distribution in the wound. Experiments on the wound treatment should be done to clarify the best spatial distribution of treatment actuators.

## Conclusion

Actuating macrophage polarization for acceleration of wound healing must be done in a narrow time interval, beginning from time of M1 maximum. Too early and too strong treatment of this type may slow down wound cleaning and lead to chronic inflammation. Too late treatment may have too small effect.

To our knowledge, this is the first work on the search of an optimal regime of wound treatment. We believe that experiments will help to clarify optimal regime for each substance. This naive modeling approach may help to predict optimal regime for the treatments with clear actions. We believe this is the first step towards smart choice of treatment regime and development of algorithms for smart wound healing devices.

## Supporting information

**S1 File. Additional results of wound healing model simulations**.

## Acknowledgments

This research is sponsored by the Defense Advanced Research Projects Agency (DARPA) through Cooperative Agreement D20AC00003 awarded by the U.S. Department of the Interior (DOI), Interior Business Center. The content of the information does not necessarily reflect the position or the policy of the Government, and no official endorsement should be inferred.

